# Functional assessment of the ionotropic receptors IR25a and IR93a in predator perception in the freshwater crustacean *Daphnia* using RNA interference

**DOI:** 10.1101/2022.02.11.480022

**Authors:** A Graeve, J Huster, J Mayweg, R Fiedler, J Plaßmann, P Wahle, LC Weiss

## Abstract

Acquiring environmental information is vital for organisms as it informs about the location of resources, mating partners, and predators. The freshwater crustacean *Daphnia* detects predator specific chemical cues released by its predators and subsequently develops defensive morphological features that reduce the predation risk. The detection of such chemical information is generally processed via distinct chemoreceptors that are located on chemoreceptor cells. Lately an ancestral type of ionotropic receptors (IRs) has been identified in crustaceans. IRs and the putative co-receptors IR25a and IR93a are postulated to be involved in chemoreception However, functional roles have not been assessed. Here, using three *Daphnia* species as model, we report that the two co-receptors are expressed within the chemosensory antennules and gene expression is increased with predator perception. Importantly, RNA interference mediated knock-down of the two IRs impedes species-specific defense expression in the three *Daphnia* species. Our results suggest that (albeit not testing the enigmatic receptor protein directly), the reduction of two associated proteins has impaired the functional aggregation of the postulated chemoreceptor complex. This in turn has hampered the perception of environmentally relevant chemical cues resulting in a substantial reduction of defensive morphological features.

## Introduction

In aquatic systems trophic interactions greatly rely on chemical signaling. Chemical cues deliver information on presence of mating partners, location of food sources and even optimal breeding grounds (Pohnert, Steinke, and Tollrian 2007; Wisenden 2000; Hay 2009; Atema 1988). Furthermore, chemical cues inform about the presence and activity of predators which in the prey can elicit defensive strategies (Tollrian and Harvell 1999). These defenses include shifts in life history parameters, alternative behavioral patterns (such as seeking refuge), or the expression of morphological defense structures imposing handling difficulties on predators. Until now, we have limited knowledge on the nature of the chemical cues (Weiss et al. 2018; Pijanowska et al. 2020; Yasumoto et al. 2005; Selander et al. 2015; Hahn et al. 2019). Even less is known about the chemoreceptive mechanisms involved in interspecific information transfer. The micro-crustacean *Daphnia* is an important component of freshwater food webs. The different *Daphnia* species display strong phenotypic plasticity against a range of predator-specific chemical cues. These cost-benefit optimized defense strategies increase organism fitness (Weiss and Tollrian 2018). For example, fatty acids conjugated to a glutamine residue are released from the phantom midge larva *Chaoborus*, and induce neckteeth expression in *D. pulex* (Krueger and Dodson, 1981; Tollrian, 1993; Weiss et al., 2016). Fish-specific bile salts (5-alpha-cyprinol) induce head- and tail-spine development in *D. lumholtzi* (Hahn & von Elert, 2022). *D. magna* reacts to fish cues by diel vertical migration (Ringelberg 2010). Cues released by the tadpole shrimp *Triops* trigger *D. magna* to grow larger and bulkier (Rabus and Laforsch 2011). Cues released from *Triops* initiate in *D. barbata* a conspicuous helmet formation and a change in body symmetry. In response to cues released by the heteropteran backswimmer *Notonecta, D. barbarta* expresses a helmet that differs in shape from that induced by *Triops* exposure (Herzog and Laforsch 2013). In *D. longicephala, Notonecta*-specific cues induce crests (Grant and Bayly, 1981; Weiss et al., 2015). This diversity of adaptive responses shows that *Daphnia* distinguish their predators based on the chemical composition of the signaling agents, and confirm the view that defense morphologies are species-specific responses to specifically impose handling problems for a selected predator.

In order to detect these sometimes highly complex compounds at extremely low concentrations (in the range of nano-or even picograms (Hahn et al. 2019; Weiss et al. 2018)) *Daphnia* need a chemoreceptive sensory system with specific chemoreceptors. Putative chemoreceptor candidates belong to a group of ligand-gated ion channels (ionotropic receptors, IRs) that evolved from ionotropic glutamate receptors (iGluRs) (reviewed in Derby et al. 2016; Benton 2015; Robertson 2015; Joseph and Carlson 2015; Zufall and Munger 2016). IRs are conserved across Protostomia including e.g. chelicerates (Rytz, Croset, and Benton 2013), myriapods (Kenny et al. 2015) and crustaceans (Eyun et al. 2017; Kozma et al. 2018; Croset et al. 2010; Derby et al. 2016). Within this class of ligand-gated ion channels, IRs are classified as tuning IRs and co-receptor IRs. Tuning IRs are ligand-specific and crucial for receptor channel function (Benton et al. 2009; Croset et al. 2010; Rytz, Croset, and Benton 2013; Eyun et al. 2017) and, given the diversity of potential ligands, presumably are highly diverse. Co-receptor IRs are co-expressed as heterotetramers associated with the tuning IRs, and are highly conserved across taxa. Co-receptor IRs have been identified in various crustaceans (Hollins et al. 2003; Corey et al. 2013; Groh et al. 2014; Zbinden et al. 2017; Groh-Lunow et al. 2015; Eyun et al. 2017). In the spiny lobster *Panulirus argus* four co-receptor IRs have been identified in the transcriptome: IR8a, IR25a, IR76b, and IR93a. The IR25a has even been shown to be functionally involved in chemoreception in the salmon louse *Lepeophtheirus salmonis*. RNAi mediated knock down of IR25a changes the louse’ sensitivity to the salmon host (Núñez-Acuña et al. 2019).

Genome data have revealed that *Daphnia pulex* has a comparatively high number of chemoreceptors among which are 58 receptors that share sequence homology to the gustatory receptor family of insects (Pe∼nalva-Arana, Lynch, and Robertson 2009). About 150 iGluR-derived IRs and three co-IRs, IR76b, IR93a and IR25a, have been identified *in silico* (Kozma, Ngo-Vu, Yan Wong, et al. 2020). However, the functional expression of these IRs and the co-IRs in *Daphnia* has not been investigated, and it is unknown whether they are functionally involved in chemoreception. Such a functional involvement could be seen in tissue-specific expression. The organ involved in predator perception was previously identified as the aesthetascs of the antennules (Weiss, Leimann, and Tollrian 2015), so that these receptors are hypothesized to be predominantly expressed in sensory organs such as the antennules. Further, the time- and environment-dependent expression changes are unknown. The functional importance of the co-IRs renders them attractive targets for experimental manipulation. Here, we studied in three *Daphnia* species (*D. longicephala, D. magna, D. lumholtzi*) the expression of the two co-IR genes *IR25a* and *IR93a*, determined the expression changes upon exposure to chemical cues released from three predators (*Notonecta* spec., *Triops* spec., three-spined stickleback *Gasterosteus aculeatus)*, and report that co-IR RNAi impairs the expression of species-specific defense morphologies.

## Results

### Co-receptor IR expression levels in the antennules, body and swimming antennae

We found a higher abundance of IR25a and IR93a mRNA in the antennules compared to the rest of the body (fig. 1 A, tab. 1). IR25a is 3.54 log2-fold higher expressed in the antennules than in the swimming antennae, while IR93a is 1.64 log2-fold higher expressed in the antennules than in the swimming antennae (fig. 1 B, tab. 1). IR25a is 4.28 log2-fold higher expressed in the antennules than in the body, while IR93a is 2.12 log2-fold higher expressed in the antennules than in the body. Both receptors are low but significantly higher expressed in the swimming antennae in comparison to the body with 0.74 log2-fold for the IR25a and 0.9 for the IR93a (fig. 1 C, tab. 1).

**Figure 1:**
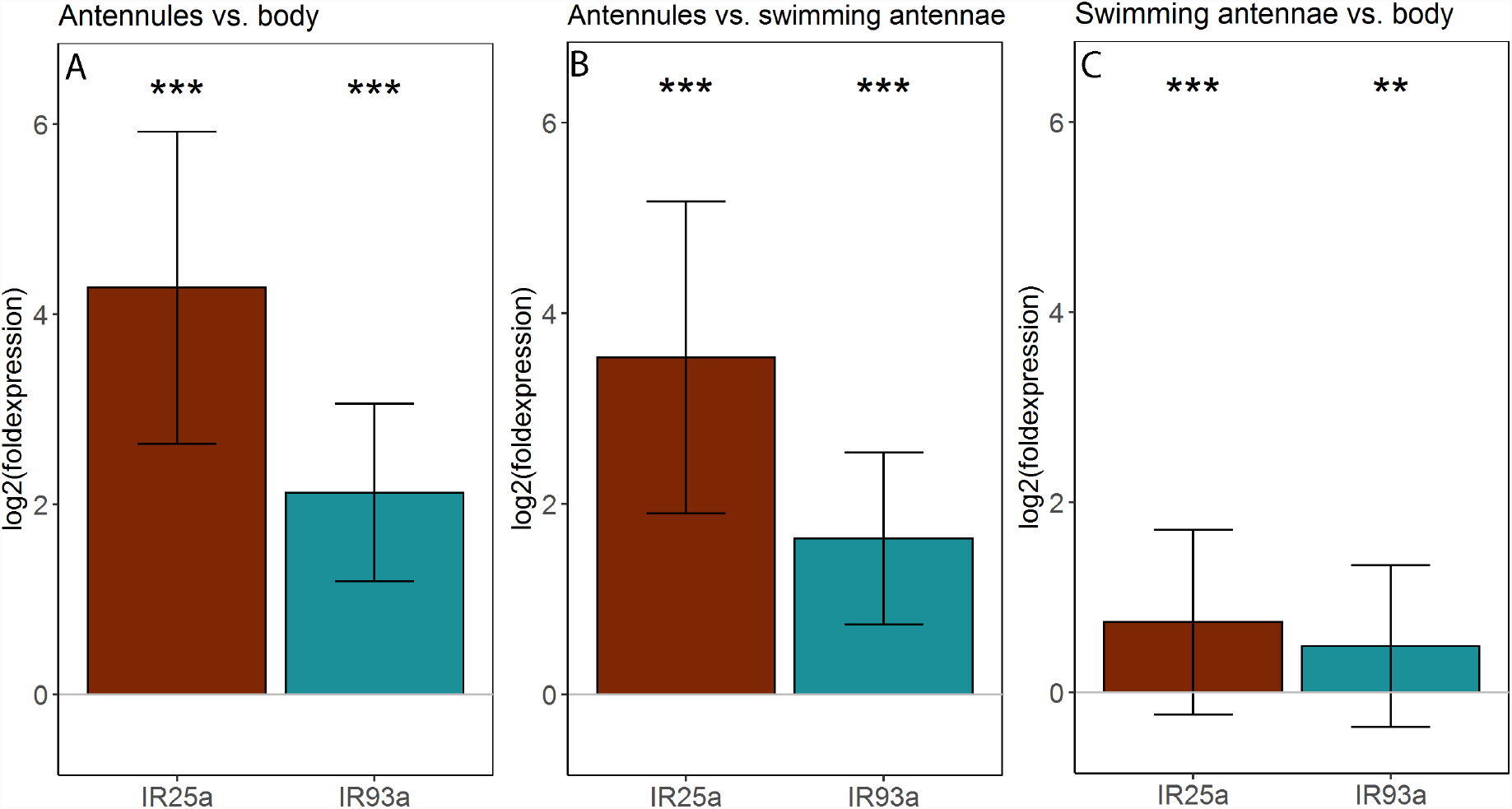
Comparative co-receptor IR expression levels in different tissues of *D. longicephala*. **A**: Increased gene expression of IR25a and IR93a in the antennules compared to the body (log2(fold expression). Both genes are significantly higher expressed in the antennules than in the body. **B:** Comparative gene expression analysis of the IR25a and the IR93a log2(fold expression) in the antennules compared to the swimming antennae. Both genes are significantly more expressed in the antennules than in the swimming antennae. **C:** Comparative gene expression analysis of the IR25a and the IR93a log2(fold expression) in the swimming antennae compared to the body. Genes are not significantly higher expressed in the swimming antennae in comparison to the body. Statistical results for one-sided T-tests are displayed in tab. 1. Bar heights depicts the mean, whiskers display ± standard deviation.

**Table 1:**
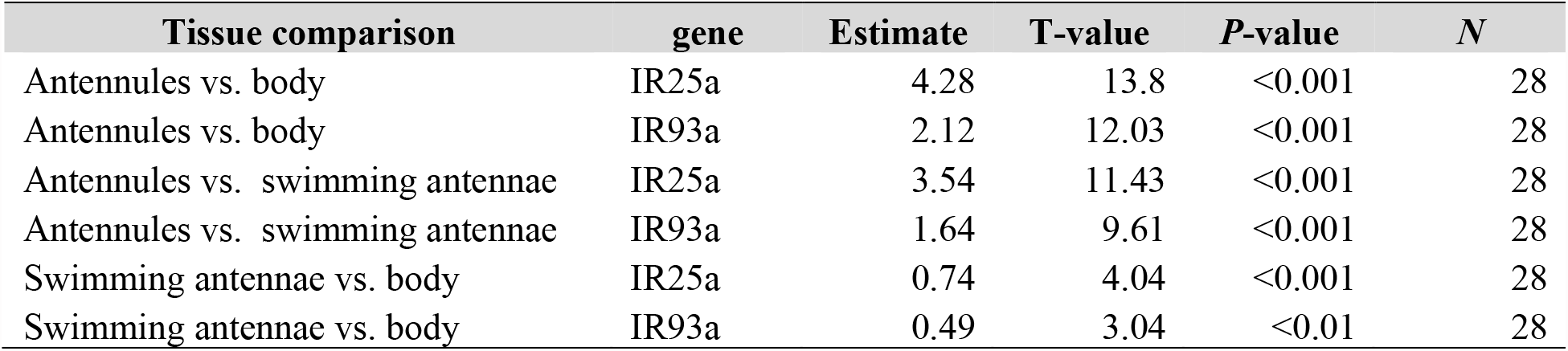
Comparative gene expression analysis of IR gene in different tissues in *D. longicephala*. T-test results of data displayed in figure 1.

| Tissue comparison | gene | Estimate | T-value | P-value | N |
| --- | --- | --- | --- | --- | --- |
| Antennules vs. body | IR25a | 4.28 | 13.8 | <0.001 | 28 |
| Antennules vs. body | IR93a | 2.12 | 12.03 | <0.001 | 28 |
| Antennules vs. swimming antennae | IR25a | 3.54 | 11.43 | <0.001 | 28 |
| Antennules vs. swimming antennae | IR93a | 1.64 | 9.61 | <0.001 | 28 |
| Swimming antennae vs. body | IR25a | 0.74 | 4.04 | <0.001 | 28 |
| Swimming antennae vs. body | IR93a | 0.49 | 3.04 | <0.01 | 28 |

### Time series of predator induced differential IR25a and IR93a gene expression

We determined the IR25a and IR93a receptors’ expression levels in a time series after predator exposure and compared it to control animals. While gene expression patterns are low and lie around 0-log2-fold during the first 4 h, it significantly increases 6 hours after predator exposure reaching a maximum expression of 1.5 log2-fold in the IR25a and 1.21 log2-fold in the IR93a. Both receptors maintain an upregulated level until 24 h (fig. 2, tab. 2).

**Figure 2:**
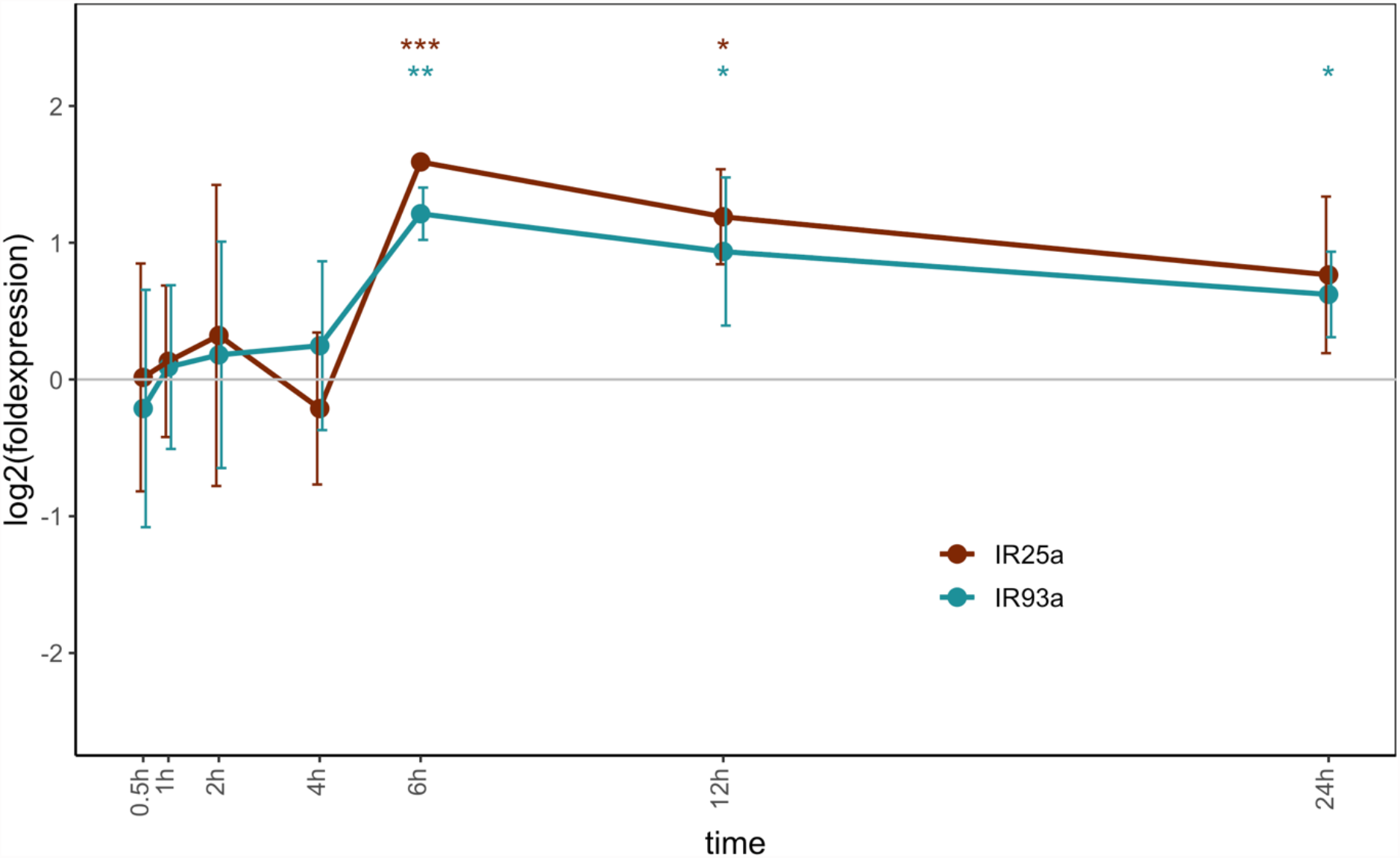
Time series of IR gene expression comparing gene expression levels of IR25a and IR93a in predator-exposed animals in comparison to control animals. IR25a gene expression is significantly up-regulated 6 h and 12 h and with a tendency at 24 h (p=0.07) post predator exposure. IR93a is significantly upregulated 6 h, 12 h and 24 h post predator exposure. Statistical results are displayed in table 2.

**Table 2:** Differential gene expression statistical analysis results comparing control and predator exposed *D. longicephala* over the measured points in time, as displayed in figure 2.

| time | gene | Mean | T-value | P-value | N |
| --- | --- | --- | --- | --- | --- |
| 0.5h | IR25a | 0.02 | 0.03 | 0.489 | 3 |
| 0.5h | IR93a | -0.21 | -0.42 | 0.643 | 3 |
| 1h | IR25a | 0.13 | 0.54 | 0.310 | 5 |
| 1h | IR93a | 0.09 | 0.34 | 0.376 | 5 |
| 2h | IR25a | 0.32 | 1.09 | 0.14 | 14 |
| 2h | IR93a | 0.17 | 0.81 | 0.21 | 14 |
| 4h | IR25a | -0.21 | -0.66 | 0.712 | 3 |
| 4h | IR93a | 0.25 | 0.69 | 0.280 | 3 |
| 6h | IR25a | 1.59 | 120.21 | <0.001 | 3 |
| 6h | IR93a | 1.21 | 10.98 | 0.004 | 3 |
| 12h | IR25a | 1.19 | 5.93 | 0.014 | 3 |
| 12h | IR93a | 0.94 | 2.99 | 0.048 | 3 |
| 24h | IR25a | 0.77 | 2.31 | 0.073 | 3 |
| 24h | IR93a | 0.62 | 3.45 | 0.037 | 3 |

### RNAi dependent knock down of predator induced morphological defenses

To study the effect of dsRNAi probes (i.e. eGFP dsIR25a, dsIR93a) we performed a full-factorial analysis of the different treatment groups (fig. 3, tab. S1, S2). eGFP-probe injected *D. longicephala* show a significantly larger relative crests height upon predator exposure in comparison to animals of the control condition. *D. longicephala* injected with dsIR25a do not show significant differences in relative crest height between the control and the predator exposed animals but are significantly smaller in comparison to the eGFP-induced group. Similarly, when injected with the dsIR93a-probe, relative crest height in predator exposed animals is not significantly different from the controls but still share similarities with the eGFP-induced group. (fig. 3 A, tab. S1, S2). This result is supported by a significant down regulation of both genes in the IR injected specimens in comparison to the eGFP injected specimens (fig. 3 B, tab. S3).

**Figure 3:**
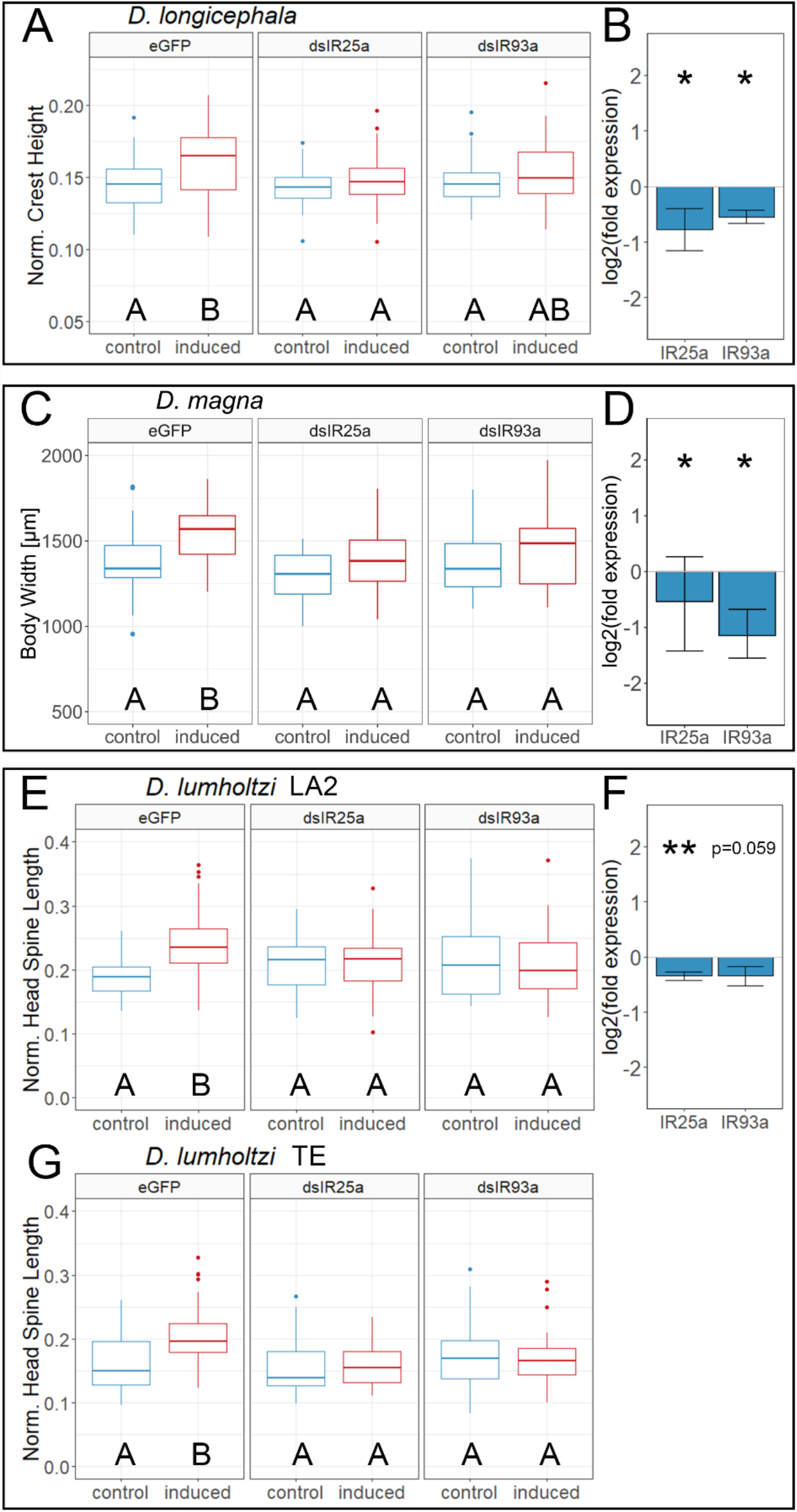
Morphological defense expression upon predator exposure after micro-injection with eGFP-, dsIR25a-or dsIR93a-probe. **A**: Normalized crest height of *D. longicephala* exposed to *Notonecta* spec. after injection with eGFP-, dsIR25a-or dsIR93a-probe. Predator exposure after eGFP-probe injection have significantly larger crests compared to the control treatment. Individuals injected with dsIR25a-or dsIR93a-probe do not express increased crest heights compared to the controls. Predator exposed dsIR93a injected *D. longicephala* injected with express crests that reach the level of the eGFP predator exposed animals. **B:** log2(fold expression) of IR25a and IR93a after probe injection. IR25a is significantly downregulated upon probe injection (log2(fold expression) = -0.77, t=-2.053, *P*=0.03). IR93a is with -0.55-log2fold significantly down regulated after probe injection, (t=-4.6, *P*=0.022). **C:** Body width of *Triops* spec exposed *D. magna* injected with eGFP-, dsIR25a-or dsIR93a-probe. Predator exposure after eGFP-probe injection induces defense significantly. Individuals injected with dsIR25a-or dsIR93a-probe do not express defensive features. **D:** log2(fold expression) of *IR25a* and *IR93a* after probe injection in *D. magna*. Both IRs are significantly downregulated upon injection with the respective probe (IR25a log2(fold expression) = -0.44, t=-2.4, *P*=0.037; IR93a: log2(fold expression) = -0.96, t=-2.79, *P*=0.025). **E:** Normalized head spine length of *D. lumholtzi* clone LA2 exposed to *Gasterosteus aculeatus* after injection with eGFP-, dsIR25a-or dsIR93a-probe. Predator exposure after eGFP-probe injection induces defenses significantly. Individuals injected with dsIR25a-or dsIR93a-probe do not express head spines. **F:** log2(fold expression) of IR25a and IR93a in *D. lumholtzi* LA2 after probe injection. IR25a is significantly downregulated (IR25a: log2(fold expression) = -0.35, t=-4.4, *P*=0.006), IR93a has a tendency to a downregulation after injection with the dsIR93a-probe (IR93a: log2(fold expression) = -0.35, t=-1.99, *P*=0.059). G: Normalized head spine length of *D. lumholtzi* clone TE exposed to *Gasterosteus aculeatus* after injection with eGFP-, dsIR25a-or dsIR93a-probe. Predator exposure after eGFP-probe injection induces significant defense expression. Individuals injected with dsIR25a-or dsIR93a-probe do not express defenses. In A, C, E, G significant differences are indicated by letters. Groups sharing the same letter are not significantly different (*P*>0.05). In B, D, E significant differences compared to the control are indicated by asterisks (\**P*≤0.05, ** *P*≤0.01 tab. S1, S2).

eGFP-probe injected *D. magna* show significantly increased body width upon predator exposure in comparison to animals of the control condition. *D. magna* injected with dsIR25a do not show significant differences in body width between the control and the predator exposed group but are significantly smaller in comparison to the eGFP-induced group. Similarly, when injected with the dsIR93a-probe, body width is not significantly different from the controls in predator exposed animals but significantly smaller in comparison to the eGFP-induced group (fig. 3 C, tab. S1, S2). This result is supported by a significant down regulation of both genes in the IR injected specimens in comparison to the eGFP injected specimens (fig. 3 D, tab. S3). Both *D. lumholtzi* LA2 *and* TE clones show a significant predator dependent increase in relative head spine length when injected with the eGFP-probe. *D. lumholtzi* injected with dsIR25a do not show significant differences in relative head spine length between the control and the predator exposed animals but have significantly smaller head spines than the eGFP-induced group. Similarly, when injected with the dsIR93a-probe, relative head spine length is not significantly different from the controls but significantly smaller when compared to the eGFP-induced group (fig.3 E, G, F tab. S1, S2). This result is supported by a significant down regulation of both genes in the IR injected specimens in comparison to the eGFP injected specimens (fig. 3 B, Stab. S3).

We repeated this experiment with alternative RNAi probes and found the same effects in the morphology of the animals (fig. 4, tab S5, S6).

**Figure 4:**
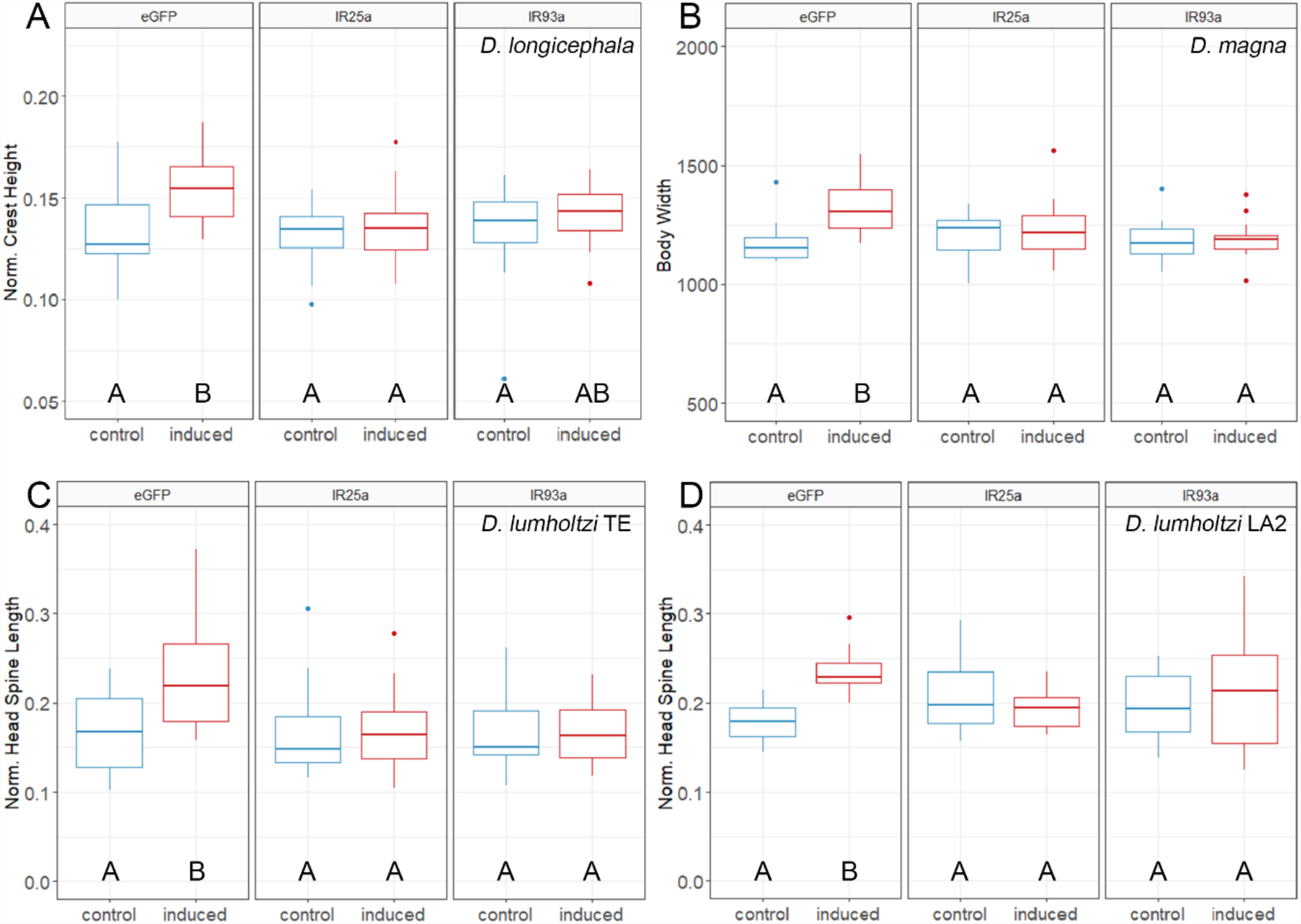
Morphological defense expression after injection with the eGFP or the alternative IR25a or IR93a dsRNA probes. A: In *D. longicephala*, normalized crest height is significantly increased upon predator exposure after eGFP probe injection. Animals injected with one of the target probes do not express crests. B: In *D. magna*, body width is significantly increased upon predator exposure after eGFP probe injection. Animals injected with one of the target probes do not express increased body width. C: In *D. lumholtzi* TE, normalized head spine length is significantly increased upon predator exposure after eGFP probe injection. Animals injected with one of the target probes do not express increased head spines. D: In *D. lumholtzi* LA2, normalized head spine length is significantly increased upon predator exposure after eGFP probe injection. Animals injected with one of the target probes do not express increased head spines. Statistical results are listed in tables S5, S6

## Discussion

Our results provide a first view on the combinatorial expression patterns of these previously *in silico* identified chemo-co-receptors in *Daphnia*.

### Co-receptor IR expression levels in the antennules

A chemosensory function of the *Daphnia* antennules has already been demonstrated through mechanical impairment of the antennules (Weiss, Leimann, and Tollrian 2015). However, the chemoreceptors expressed by the antennules remained undetermined. We here report a higher abundance of IR25a and IR93a mRNA in the antennules compared to the rest of the body. In the swimming antennae, we observe a somewhat higher (∼0.5 log2-fold) level of both co-receptor IRs in comparison to the body. Based on these results, we also expect to find more receptor protein molecules in the antennules than in swimming antennae and even less in the body.

Comparing the degree of the expression levels of both receptors, the IR25a seems to be more abundant in the antennules than the IR93a. In the swimming antennae both receptors seem equally expressed. This further supports the idea that the antennules serve as a functional unit of chemoreception. In different crustacean taxa, a high RNA expression level of the IR25a coincides with the location of the IR25a protein on the lateral flagella (bearing the aesthetascs) that mediate chemoreception (Tadesse et al. 2011; Stepanyan et al. 2004; Kozma, Ngo-Vu, Rump, et al. 2020; Kozma et al. 2018; Kozma, Ngo-Vu, Yan Wong, et al. 2020; Corey et al. 2013; Groh-Lunow et al. 2015; Zbinden et al. 2017; Hollins et al. 2003). This implies that also in the onchyuran *Daphnia* the IR25a and IR93a proteins are localized on the antennules and potentially engage in the perception of chemical cues. To further support our observations, antibody staining directed against IR25a and IR93a would be necessary. Unfortunately, the only antibody that is currently available (product #EKY005, Kerafast USA), is directed against the IR25a in lobster and gave unreliable staining patterns and Western Blot validation failed; a similar effect was observed in salmon lice (Komisarczuk, Grotmol, and Nilsen 2017).

While we now have strong evidence that both co-IR proteins are located on the antennules, we still know little about their specific localization in the body and the swimming antennae. In case there are peripheral receptors expressed, these could either also be of chemosensory type, or they could be thermo-sensitive as has been demonstrated when the IR25a or IR93a are co-expressed with e.g. the IR21a in *Drosophila* (Ni 2021). These receptor classifications will be tasks for future investigations.

As chemoreceptors are often subject to an increased turnover rate when being stimulated with their ligands (Koerte et al. 2018), we investigated the effect of predatory chemical cues on differential gene expression in a time dependent manner.

### Time dependent gene expression patterns of IR25a and IR93a

We compared expression levels in the antennules of predator exposed to control *D. longicephala*. Six hours post predator exposure we detect a significant upregulation of both IRs. Together with the subsequent continuous upregulation of both receptors, this probably marks the onset of an increased receptor turn-over rate which is due to ligand-receptor binding activity (Koerte et al. 2018). It is important to mention that this does not depict the point in time for the onset of predator perception. This must lie earlier, if not even instantaneous, as increased cell proliferation patterns have been reported already two hours post predator exposure (Graeve et al. under revision). Ecologically, predator detection resulting in the fast onset of the mechanisms engaged in defense expression is critical. The longer the time lag between predator perception and the onset of such mechanisms, the longer it takes for the defenses to serve as an effective protection (Clark and Harvell 1992; Jeschke, Laforsch, and Tollrian 2008; Hoverman and Relyea 2007; Gabriel et al. 2005).

In *Drosophila* such a ligand-dependent up-regulation in the IRs’ mRNA expression levels has not been reported (Koerte et al. 2018) and we therefore speculate differences in the receptors’ relevance for detecting chemical agents in these two taxa. *Drosophila* possesses a highly divergent group of ORs whose mRNA expression levels change in response to ligands, but these ORs are absent in *Daphnia*. Together with the findings reported by Kozma et al. 2020, we hypothesized, that chemoreception in *Daphnia* is a central function of the IRs. To support this idea, we tested the functional involvement of these co-IRs using RNAi.

### Functional analysis of IR25a and IR93a

All results were reproducible with the alternative probes, targeting a different gene region. Together with qPCR results, we detect a successful knock-down of our target genes in all species. Injection of the eGFP control probe does not affect defense expression in any of the tested species. However, when injected with the IR25a probe, the three *Daphnia* species lose their ability to react to predators. All measured parameters are not significantly different from the control animals but significantly different from the predator exposed animals, so that defenses are not at all expressed. When injected with the IR93a probe defenses in *D. magna* and the two *D. lumholtzi* clones, are again not at all expressed. Only, in *D. longicephala* defenses are significantly reduced but do not reach the control level, indicating a smaller involvement of the IR93a in the perception of *Notonecta* specific signaling cues.

This shows that both IRs are functionally involved in the perception of the here involved predator specific chemical cues. Our observations could be explained by two scenarios 1.) The IR25a and the IR93a assemble and function as one chemoreceptor unit. However, as this receptor combination was shown to function as a cool sensor and detected in the periphery of the chemosensory organs in *Drosophila*, we anticipate it to be unlikely to have chemosensory abilities in *Daphnia* (Knecht et al. 2016). 2.). Each of the two IRs function as chemo-co-receptors and assemble with respective tuning receptors to form functional chemoreceptor units. This also explains why the chemosensory capacities i.e. in detecting and reacting to this range of chemosensory signals are hampered, as the knock-down of both IRs impairs receptor assembly and trafficking. Even though the chemical composition of the *Triops* and the *Notonecta* specific signaling cues have not been elucidated, we are highly confident that they are different in their nature, due to their phylogenetic distance and the distinct responses they induce in the different prey species (reviewed in Weiss & Tollrian 2018). Further, bile salts are specific to vertebrates and most likely not present in any invertebrate taxon (Hahn et al. 2020). Together with the fact, that even the experimentally necessary environmental transfer induced the differential expression of both receptors further supports their co-receptor function. Interesting is that the knock-down of the IR93a in *D. longicephala* only reduces but not prevents defense expression. This indicates that the perception of the cues associated with the *Notonecta* kairomone are predominantly perceived via IR25a receptor functioning and to a less extent via the IR93a. Future studies will aim to identify the concrete tuning IR(s) that assemble with the here identified co-IRs mediating chemoreception of natural substances in the environment. This also lays the fundamental basis for understanding how *Daphnia* is able to react to changes in the chemical environment and adapt through phenotypic plasticity. With respect to sensory ecology, knowing the involved receptors may also guide to disentangling chemical signaling agents, whose chemical identification is often the search for the needle in the haystack. By this we will be able to better understand organism-by-environment interactions and how this can have an ecosystem-wide effect.

## Material and Methods

### *Daphnia* cultures

*Daphnia longicephala* clone LP1 (Lara-Pond, Australia) and *D. lumholtzi* clones LA2 (Louisiana, USA; kindly provided by Ramcharan) and TE (Fairfield Reservoir, Texas, USA; kindly provided by Sterner) were cultured in artificial *Daphnia* medium (ADaM (Klüttgen et al. 1994)) in 1 L beakers (Weck®, Germany) containing 20 - 25 age-synchronized individuals. *D. magna* clone FT44-2 (Rockpool 44, Tvärminne, Finland, kindly provided by D. Ebert) was cultured in charcoal-filtered tap water (to cohere with the culture conditions of *Triops*) in 1 L beakers containing 10 to 15 age-synchronized individuals. Animals were kept under constant day:night conditions (16 h:8 h) at 20 ± 1°C and fed *ad libitum* with the algae *Acutodesmus obliquus*. The beakers were cleaned every 48 hours to remove exuviae, debris and excess algae. Half of the medium was exchanged weekly.

### Predator cultures

Backswimmers (*Notonecta* spec.) were collected from the ponds of the Ruhr-University Botanical Garden. Animals were maintained in 1 L beakers (WECK^®^, Germany) filled with charcoal-filtered tap water under standardized conditions (20 ± 1°C with 16 h:8 h light:dark cycle). They were fed *ad libitum* with *Daphnia* spec.. *Triops* spec. were raised from sand containing the eggs. Animals were kept in 1 L plastic tanks filled with one part charcoal-filtered tap water and one part pre-desalted water under standardized conditions (20 ± 1°C with 16 h:8 h light:dark cycle). *Triops* spec. were fed daily with fish flakes (Futter Granulat, MultiFit) (flakes were crushed for freshly hatched animals) and after reaching a body length of ∼0.5 cm were fed additionally with *Daphnia* spec..

Three-spined sticklebacks (*Gasterosteus aculeatus*) (Zoo Zajac, Duisburg, Germany) were cultured in ADaM in a 60 L glass tank at a constant temperature of 20 ± 1°C with 16 h:8 h light:dark cycle. Fish were fed *ad libitum* every 24 h with living *Daphnia* spec. and frosted *Chironomus* larvae (Amtra, Germany). All animals were kept under conditions complying with care and welfare.

### RNAi-probe design and synthesis

RNAi-probes were designed against transcriptomic data of *D. longicephala* and *D. lumholtzi*, and the online deposited *D. magna* genome data (https://metazoa.ensembl.org/Daphnia_magna/Info/Index). Probes were cloned in-house following the requirements and strategy published by (Knorr et al., 2013; Posnien et al., 2009,, fig. 5). For reverse transcription of a ∼500 bp probe, we designed primers in Primer3 (tab. 3): Length: 19 – 22 bp; Max Self Complementary: 5; Max 3’ Self Complementary: 1; Max Pair End Complementary: 1. The desired annealing temperature was set to 60 °C. BLAST homology search was performed for the whole probe sequence with settings optimized for ’somewhat similar sequences’ (blastn) to exclude off-targets.

**Figure 5:**
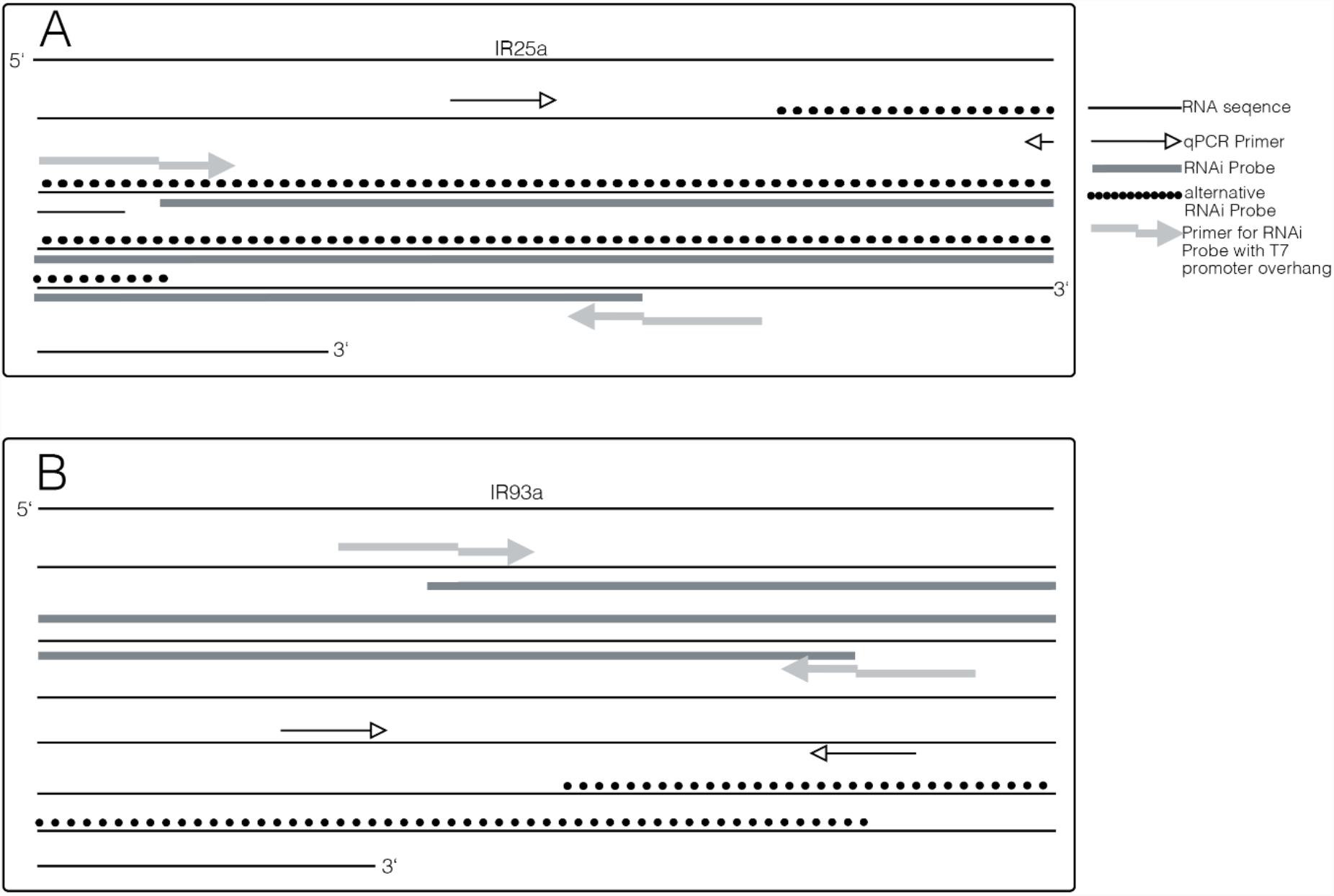
Probe design. Extract of the IR25a (A) consensus and IR93a (B) obtained from the sequences of *D. longicephala, D. magna* and *D. lumholtzi*. Displayed are the positions of the qPCR primers (black arrow), the primers for our custom designed RNAi probe (grey arrow) and the respective probe sequence (grey line) and the universal RNAi probe sequence (dotted line).

**Tab 3:**
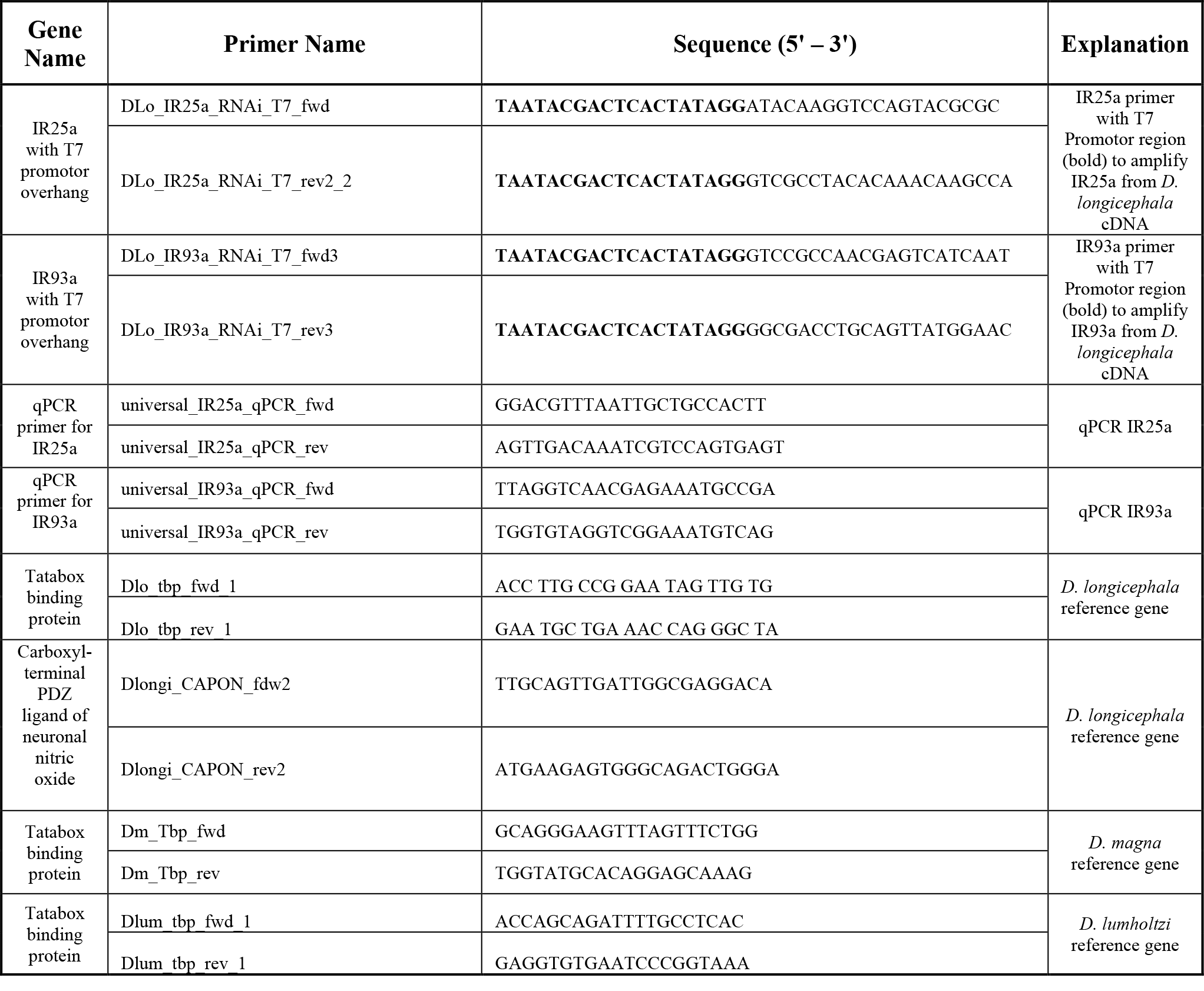
List of primers used to synthesize RNAi probes from cDNA and qPCR

For subsequent *in vitro* transcription with T7 polymerase, T7 promotors were added via PCR using GoTaq Master Mix (Promega) on *D. longicephala* cDNA. The obtained amplicon was used for TA-cloning into a pCRII vector (TOPO TA Cloning kit, Invitrogen) and transformed into TOP 10 *E. coli* (Invitrogen). After Midi-prep (Nucleo Bonda Xtra Column Filter kit, Machery & Nagel) samples were Sanger-sequenced for validation. *In vitro* transcription was conducted using the HiScribe T7 High Yield RNA Synthesis kit (New England Biolabs). Finally, double-stranded RNA-probes (dsRNAi-probes) were cleaned using the Monarch RNA clean-up kit (New England Biolabs, Germany).

As a negative control, we purchased an eGFP-probe as *Daphnia* do not have this sequence encoded in the genome (Eupheria Biotech, Dresden, Germany; Sigma Product ID: EHUEGFP). We validated our findings using probes that lie in a different region of the IR25a and IR93a gene. In the case of the IR93a this alternative probe targets a region that does not overlap with our initial probe. The IR25a sequence did not permit such an alternative non-overlapping probe, and therefore, we generated a probe that overlaps with the other probe by 426 bp. These control probes were also purchased from Eupheria Biotech (Dresden, Germany). All RNAi-probes were diluted to a final concentration of 300 ng/µL in 4 µL aliquots and stored at -80 °C until use. We further tested for sequence homology of the IR25a probe to the IR93a gene and vice versa, yielding only weak homology. Sequence homology matrix between probes and target sequence are given in table 4.

**Tab 4:**
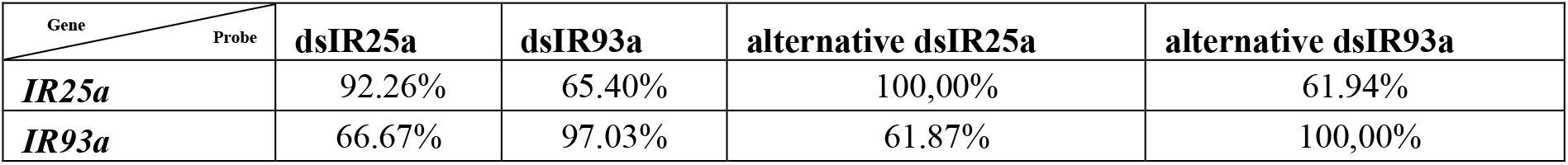
Sequence homology between *IR25a* and *IR93a* consensus sequences obtained from *D. longicephala, D. magna* and *D. lumholtzi* and injected RNAi probes. dsIR25a and dsIR93a probes are designed with *D. longicephala* cDNA. The alternative RNAi probes targeting different sequence regions of the genes “universal dsIR25a” and “universal dsIR93a” were designed for the consensus sequence of all three species. Alignments were performed using the EMBL-EBI Multiple Sequence Alignment tool (Madeira et al. 2019).

### Predator exposure bioassays

*D. longicephala* were exposed to *Notonecta* spec.. For that, we used a 12 L glass tank filled with 4 L ADaM. Seven net cages (mesh size 150 µm), each containing one Notonectid were placed into the tank. Predators were fed 10 adult *Daphnia* each. After 24 hours two Notonectids were removed to avoid over-exposure and to create space for net cages with *D. longicephala*. 3-day old *D. longicephala* were added to the tanks in net cages containing a maximum of 20 *Daphnia* and fed *ad libitum* with *Acutodesmus obliquus* algae.

*D. magna* were exposed to *Triops* spec. in 12 L glass tanks filled with 3 L charcoal-filtered tap water, a layer of approximately 0.5 cm sand and three *Triops* spec. (with <1 cm body length). Predators were fed *ad libitum* with 3-day old *D. magna* and fish flakes. After 24 hours net cages with 4-day old test specimens were added to the tank and fed *ad libitum* with *Acutodesmus obliquus*.

To also exclude that are effects are of clonal origin, we also tested two clones of the same species. We exposed *D. lumholtzi* clones TE and LA2 to *Gasterosteus aculeatus* in12 L glass tanks filled with 8 L ADaM and two sticklebacks with a body length of 3 – 5 cm. After 24 h net cages with 3-day old *D. lumholtzi* were added to the tank and fed *ad libitum* with *Acutodesmus obliquus* algae.

### Controls for predator bioassays

All controls were handling controls conducted at the same time with the same set of equipment and liquid batches lacking just one variable: no predators were added to the tanks.

### Tissue specific sampling to determine IR25a and IR93a localization

To determine co-receptor localization, we dissected three-day old *D. longicephala* in a tissue-specific manner, removing the antennules (by cutting off the rostral tip), and swimming antennae, which were dissected with metal tweezers. Antennules, swimming antennae and the body were then transferred into tubes containing DNA/RNA shield (Zymo Research). Samples were directly manually homogenized with plastic pistils and stored at -80 °C until further usage. Each biological replicate contained 40-60 animals which we found to provide enough RNA for quantitative PCR.

### Predator-induced differential gene expression levels of IR25a and IR93a in the antennules

To investigate predator-induced differential IR gene expression, we exposed 40-60 three-day old *D. longicephala* to *Notonecta* spec. as described in the induction assay for 0.5 h, 1 h, 2 h, 4 h, 6 h, 12 h, and 24 h. Subsequently, animals were killed, and antennules were cut off with a sharp razor blade. Again, samples were homogenized and stored in DNA/RNA Shield™ (Zymo Research, Germany) at -80 °C until further usage. Each biological replicate contained 40-60 animals providing good quality and quantity yields for gene expression analysis.

### Quantitative reverse transcription PCR (RT-qPCR)

RNA for RT-qPCR was extracted with the Quick-RNA Microprep Kit (Zymo Research) and diluted to a working concentration of 10 ng/µL. The Luna® One-Step RT qPCR Kit (New England Biolabs, Germany) was used for RT-qPCR in the LightCycler® 96 System (Roche, Germany) on 96 well plates (MJ white, Sarstedt, Germany) according to the protocol with a modified total reaction volume of 10 µL. For each treatment, a minimum of three biological and two technical replicates (from which we calculated the mean CT) were used. Primer efficiency was calculated using LinReg PCR (Ruijter et al. 2009). All primers had an efficiency of >0.87<1.0. We analyzed log2-fold expression changes based on the ddCT method (Livak and Schmittgen 2001) with respect to primer efficiency using specific reference genes chosen based on geNorm (Vandesompele et al. 2002) (tab. 5). Differential expression was determined with a one-sample T-test on the log2-fold expression data.

**Tab 5:**
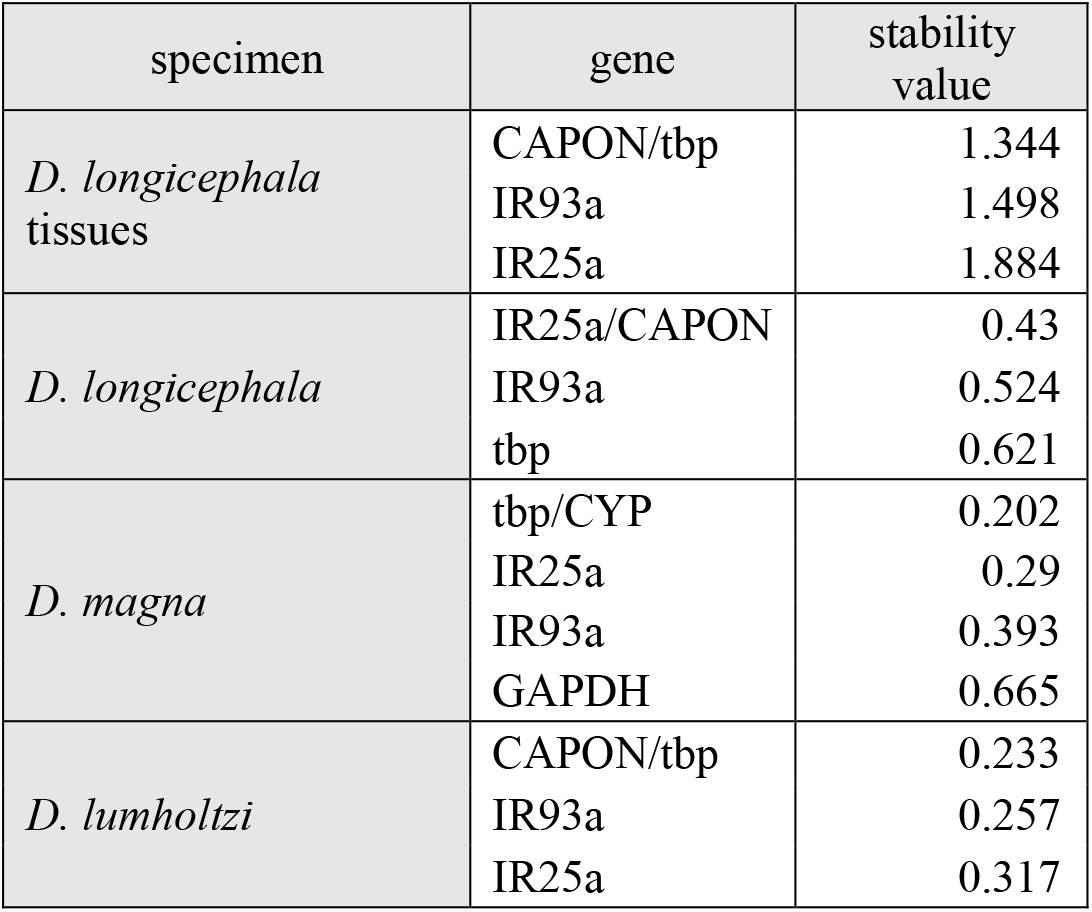
Validation of reference genes. **A:** Gene stability in *D. longicephala* antennules, 2^nd^ antennae and body. Data of induced and control and injected and non-injected samples from all observed sampling points were used. **B:** Gene stability in *D. longicephala*. Data of injected (eGFP, dsIR25a or dsIR93a) and non-injected samples were used. **C:** Gene stability in *D. magna*. Data of injected (eGFP, dsIR25a or dsIR93a) and non-injected samples were used. **D:** Gene stability in *D. lumholtzi*. Data of injected (eGFP, dsIR25a or dsIR93a) and non-injected samples were used.

### Microinjection

For microinjections we used the FemtoJet 4i (Eppendorf) attached to a micromanipulator (Narishige). Capillaries (without filament, length: 100 mm, outer diameter: 1 mm, inner diameter: 0.58 mm, Hilgenberg, GmbH, Germany) were pulled using the Micropipette Puller P-1000 (Sutter Instruments) with the following settings: ramp = 505, heat = 500, pull = 40, velocity = 60, time = 120, pressure = 500. Phenol red was added in a ratio of 1:5 to the RNAi-probes for visual control of injection success. Probes were filled into the capillary using a fine glass pipet that was pulled out over an open flame. Filled injection capillaries were opened by breaking the tips with tweezers and sharpened over a wet stone. Just prior to injection the probes (300 ng/µl) were diluted with phenol red using a ratio of 1:5 to obtain visual control over the injection success. We validated that phenol red and eGFP do not affect *Daphnia* viability or their ability to express inducible defenses (fig. 6).

**Figure 6:**
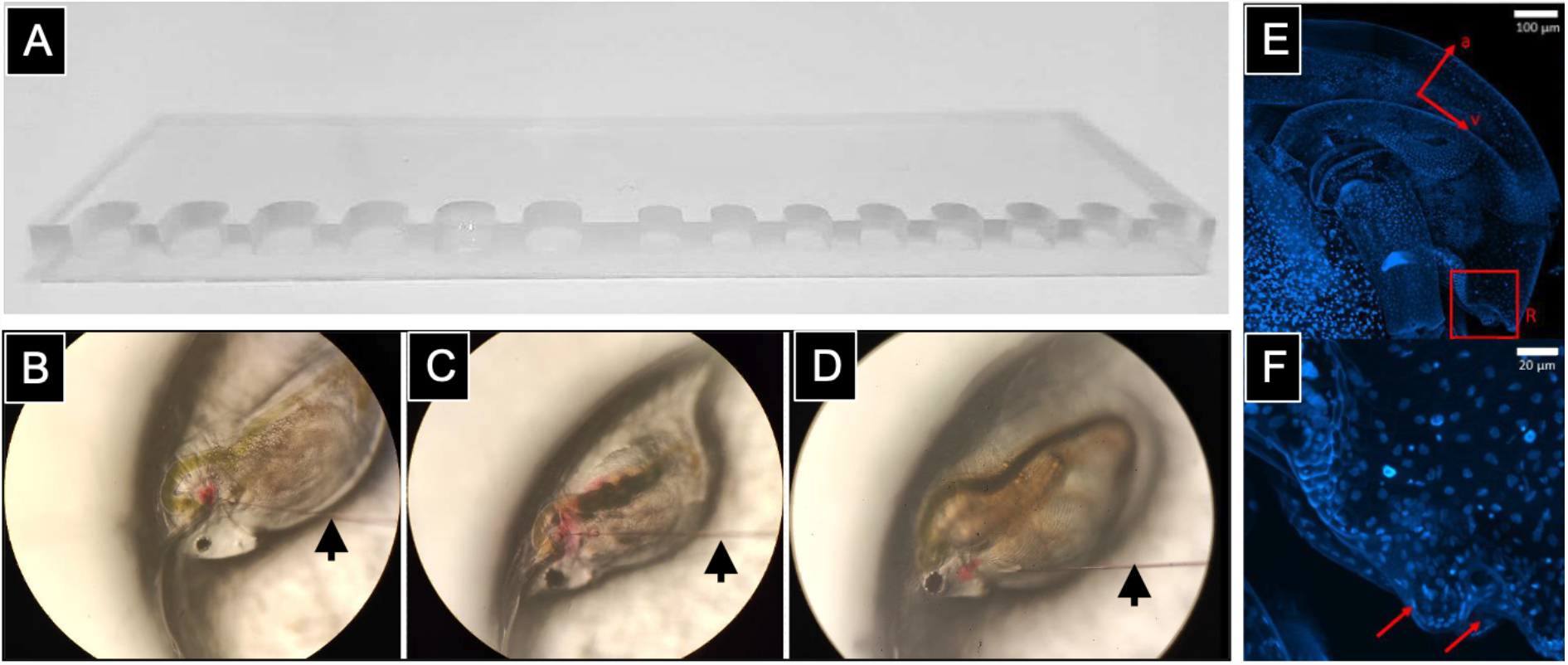
Methodology used to inject RNAi probes in juvenile *Daphnia* **A**: custom designed object slide holder with U-shaped notches to position the different *Daphnia* species during microinjections. **B**: Example for microinjections performed with *D. longicephala*, using phenol red as a visual guidance of successful injection **C**: Example for microinjections performed with *D. lumholtzi*, and **D**: Example for microinjections performed with *D. magna*. **B-C**: Arrows mark the injection needle. **E, F**: Display of Hoechst injected *D. magna*, where the microinjection of the cell marker dye, successfully reaches and stains, cells of the head and especially the basis of the antennules. Boxes mark the region of the antennules, red arrows depict sensory somata at antennules’ base.

For the injection of the RNAi probes, we established a microinjection setup in which we injected juvenile and adult *Daphnia* of the different species. For this we designed a Plexiglas object slide (8 cm x 2.5 cm x 0.4 cm), containing u-shaped notches of 0.5 cm and 0.25 cm diameter to fit the respective sizes of different *Daphnia* age and size classes (fig. 6 A). For injection the *Daphnia* were held in place by placing the *Daphnia* backwards against the frame of the notch (fig. 6 B, C, D). The injection needle was positioned on the head near the base of the 2^nd^ antenna (fig. 6 B, C, D). Injection pressure was set to 300 hPa, holding pressure was 30 hPa. *D. longicephala* were injected in the neck region near the base of the swimming antenna, while *D. magna* and *D. lumholtzi* were injected through the underside of the rostrum. The injection lasted 3-12 s until a red stain inside the *Daphnia* indicated the injection success. Subsequently, the viable and injected *Daphnia* were transferred into snap-cap vials containing 100 ml ADaM, or kairomone enriched ADaM, respectively. We ensured that injected media can also reach the cells of interest by injecting Hoechst as nuclear marker. This stained all body cells including the cells at the base of the antennules (fig. 6 E, F).

### Functional assessment of IR25a and IR93a

To study the function of the two putative co-receptors, we microinjected three-day old *D. longicephala*, four-day old *D. magna* and three-day old *D. lumholtzi* (clone TE and LA2) with the dsRNAi-probes. After 2 to 3 hours the animals were transferred to the respective predator treatment. The phenotypes were then analyzed in the instar where defense expression normally occurs, which is usually two molting cycles after predator exposure, so approximately 48 hours post predator exposure (fig. 7). We validated down-regulation of the target genes again using qPCR. For this we collected 5*10 *D. longicephala*, 5*10 *D. lumholtzi* and 5*5 *D. magna* 4 to 6 h post microinjection in RNA shield.

**Figure 7:**
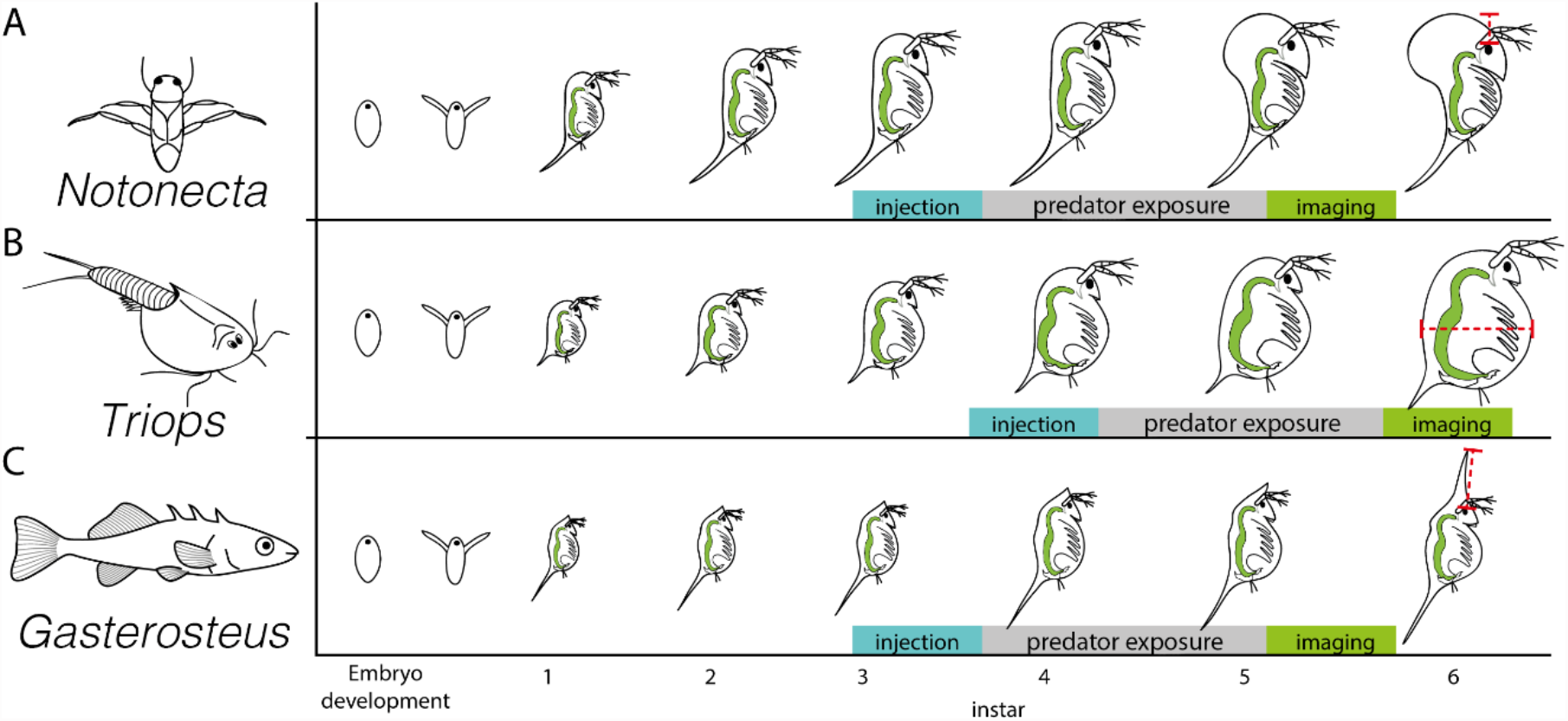
Schematic display of the experimental strategy used for RNAi. **A:** *D. longicephala* was microinjected with the respective RNAi probes once the animals reached the third instar. After three hours the animals were transferred to the predator *Notonecta* or control treatment respectively. Defense expression was measured as crest height in the fifth instar. **B:** *D. magna* was injected with RNAi probes in the fourth instar. Three hours after microinjection, the animals were transferred to the predator *Triops* or the control condition respectively. Defense expression was measured as body width in the sixth instar. **C:** Both *D. lumholtzi* clones LA2 and TE, were microinjected with both the respective RNAi probes in the third juvenile instar and exposed to the predator *Gasterosteus aculeatus* two to three hours later. Defense expression was measured as head spine length in the fifth instar.

### Morphological defense measurement

Crest height in *D. longicephala* was measured in a vertical line from the upper margin of the compound eye to the most distal point of the head. As this trait is correlated with the body size, we normalized crest height by dividing crest height by body length. Body length was measured from the upper margin of the compound eye to the root of the tail spine. Body width in *D. magna* was measured in a horizontal line at the widest point of the body. Head spine length in *D. lumholtzi* was measured from the upper margin of the compound eye to the tip of the head spine and again normalized by body length as measured above (red lines in fig. 7). For measurements we used an image analysis system composed of Olympus SZX16 stereo microscope equipped with a digital camera (Olympus DP74) controlled by the software cell sense. Measurements were taken on the images using the line tools.

### Data management and statistical analyses of morphological defenses and RNAi probe injection effects

All data was collected in Excel (Microsoft Inc.). We tested for normal distribution of the data using a Shapiro-Wilk test. As the data for morphological defense expression after probe injection did not follow a normal distribution, we log-transformed the data, which was then normally distributed. To determine the effect of the injected probes, we used an analysis of variance (ANOVA) with injected probe and treatment as factors. To determine differences between individual groups analysis of variance was followed by a Bonferroni post hoc test. Data is displayed untransformed. Statistical analyses and plots were made in R using the ggplot2 package (R Development Core Team 2011; Hadley 2016).

## Supplementary Material

**Figure S1:**
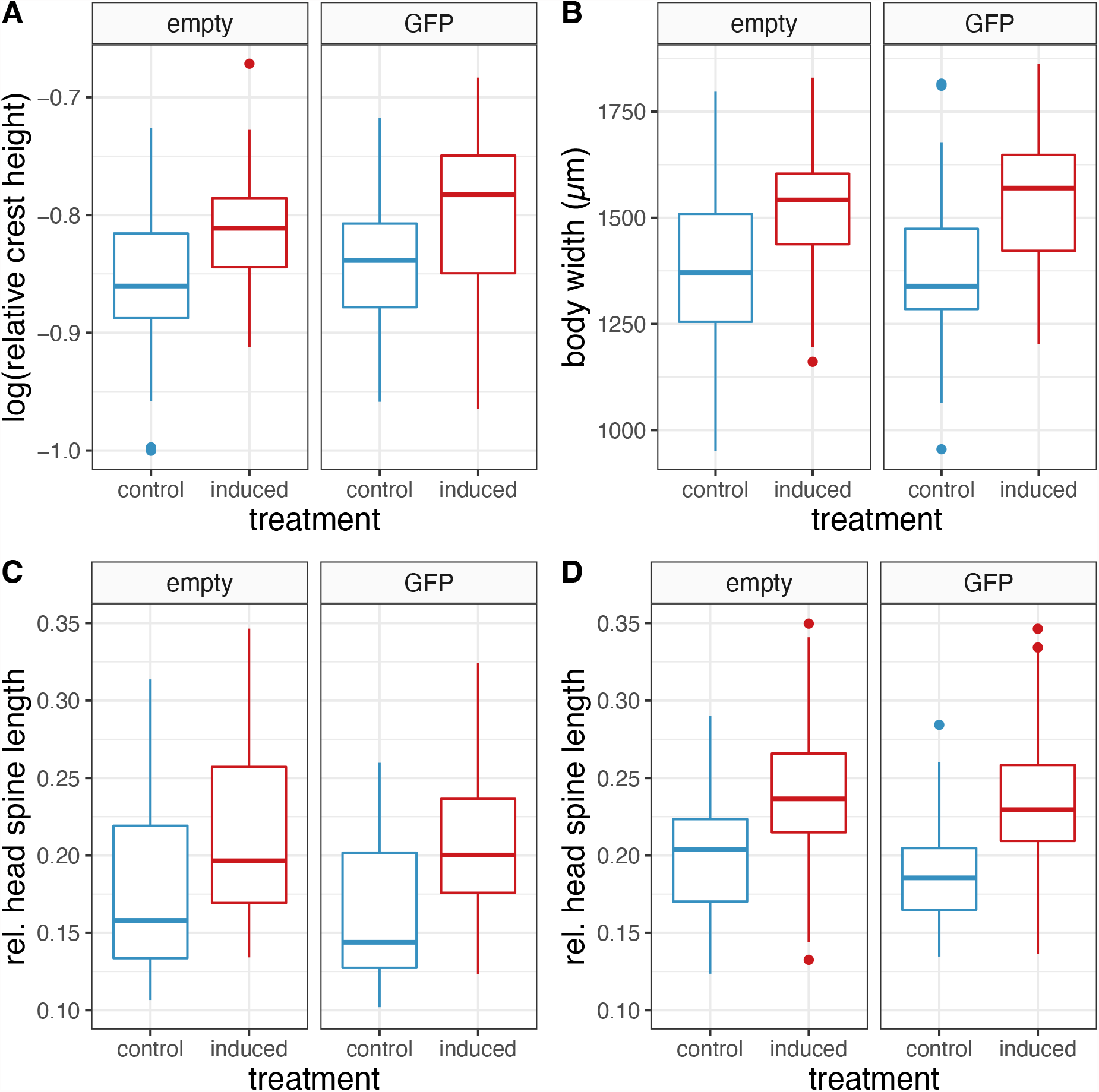
**e**GFP dsRNA probe injection does not affect the expression of morphological defenses in **A:** *D. longicephala* significantly expresses crests: ANOVA_treatment_ F(1, 0.11)=37.490; p<0.001, which is unaffected by the dsGFP RNA probe injection ANOVA_injection_ F(1, 0.01)=3.281; p0=0.07 **B:** *D. magna* significantly expresses deeper bodies: ANOVA_treatment_ F(1, 176229)=79.233; p<0.001, which is not affected by dsGFP probe injection ANOVA_injection_ F(1, 4327)=0.195, p=0.659 **C:** *D. lumholtzi* TE significantly expresses head spines: ANOVA_treatment_ F(1, 0.11894)=38.112; p<0.001, which is unaffected by dsGFP RNA probe injection: ANOVA_injection_ F(1, 0.00791)=2.534, p=0.113 and **D:** *D. lumholtzi* LA2 significantly expresses head spines: ANOVA_treatment_ F(1, 0.15336)=78.405; p<0.001, which is unaffected by dsGFP probe injection ANOVA_injection_ F(1, 0.0048)=2.46, p=0.118

## Statistical results of RNAi knock down

**Table S1:** Factorial ANOVA on log(normalized crest height) in *D. longicephala*, log(body width) in *D. magna* and log(normalized head spine length) in *D. lumholtzi* clones TE and LA2 after injection with eGFP-, dsIR25a-or dsIR93a-probe measured after 48 hours predator exposure or control treatment. Related to fig. 5. Abbreviations: SS, square sum; MS, mean square.

|  | SS | Degree of Freedom | MS | F | P | Partial eta-square | Non-centrality | Observed power |
| --- | --- | --- | --- | --- | --- | --- | --- | --- |
| *Daphnia longicephala* |  |  |  |  |  |  |  |  |
| Intercept | 219.66 | 1 | 219.66 | 81025.21 | ≤0.001 | 1.00 | 81025.21 | 1.00 |
| Injection | 0.01 | 2 | 0.01 | 2.74 | 0.066 | 0.02 | 5.49 | 0.54 |
| Treatment | 0.05 | 1 | 0.05 | 16.78 | ≤0.001 | 0.05 | 16.78 | 0.98 |
| Injection*Treatment | 0.02 | 2 | 0.01 | 3.56 | 0.029 | 0.02 | 7.13 | 0.66 |
| Error | 0.86 | 317 | 0.00 |  |  |  |  |  |
| *Daphnia magna* |  |  |  |  |  |  |  |  |
| Intercept | 3027.82 | 1 | 3027.82 | 1482460.91 | ≤0.001 | 1.00 | 1482460.91 | 1.00 |
| Injection | 0.05 | 2 | 0.03 | 13.13 | ≤0.001 | 0.07 | 26.25 | 1.00 |
| Treatment | 0.06 | 1 | 0.06 | 31.46 | ≤0.001 | 0.09 | 31.46 | 1.00 |
| Injection*Treatment | 0.02 | 2 | 0.01 | 5.87 | 0.003 | 0.03 | 11.73 | 0.87 |
| Error | 0.690 | 338 | 0.00 |  |  |  |  |  |
| *Daphnia lumholtzi* clone TE |  |  |  |  |  |  |  |  |
| Intercept | 150.76 | 1 | 150.76 | 12870.64 | ≤0.001 | 0.98 | 12870.64 | 1.00 |
| Injection | 0.21 | 2 | 0.10 | 8.76 | ≤0.001 | 0.07 | 17.53 | 0.97 |
| Treatment | 0.11 | 1 | 0.11 | 9.36 | 0.002 | 0.04 | 9.36 | 0.86 |
| Injection*Treatment | 0.16 | 2 | 0.08 | 6.80 | ≤0.001 | 0.05 | 13.61 | 0.92 |
| Error | 2.823 | 241 | 0.01 |  |  |  |  |  |
| *Daphnia lumholtzi* clone LA2 |  |  |  |  |  |  |  |  |
| Intercept | 109.99 | 1 | 109.99 | 13966.74 | ≤0.001 | 0.98 | 13966.74 | 1.00 |
| Injection | 0.01 | 2 | 0.01 | 0.65 | 0.523 | 0.01 | 1.30 | 0.16 |
| Treatment | 0.09 | 1 | 0.09 | 11.57 | ≤0.001 | 0.05 | 11.57 | 0.92 |
| Injection*Treatment | 0.15 | 2 | 0.08 | 9.70 | ≤0.001 | 0.07 | 19.39 | 0.98 |
| Error | 1.929 | 245 | 0.01 |  |  |  |  |  |

**Table S2:**
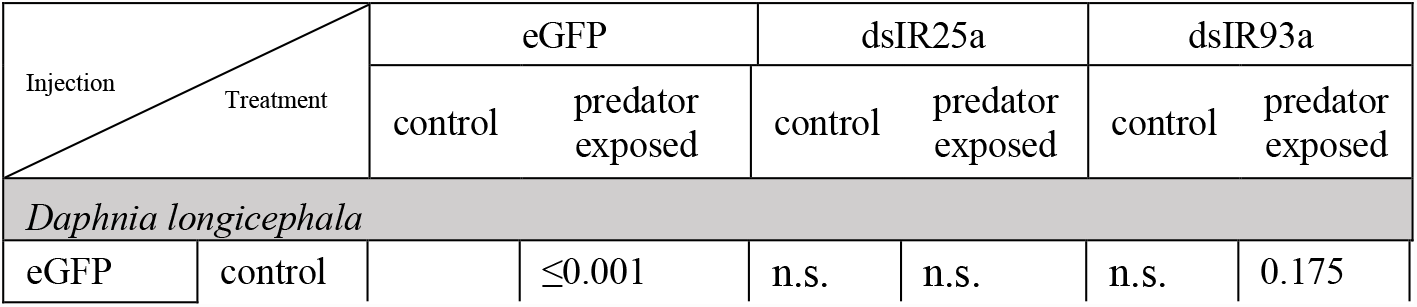
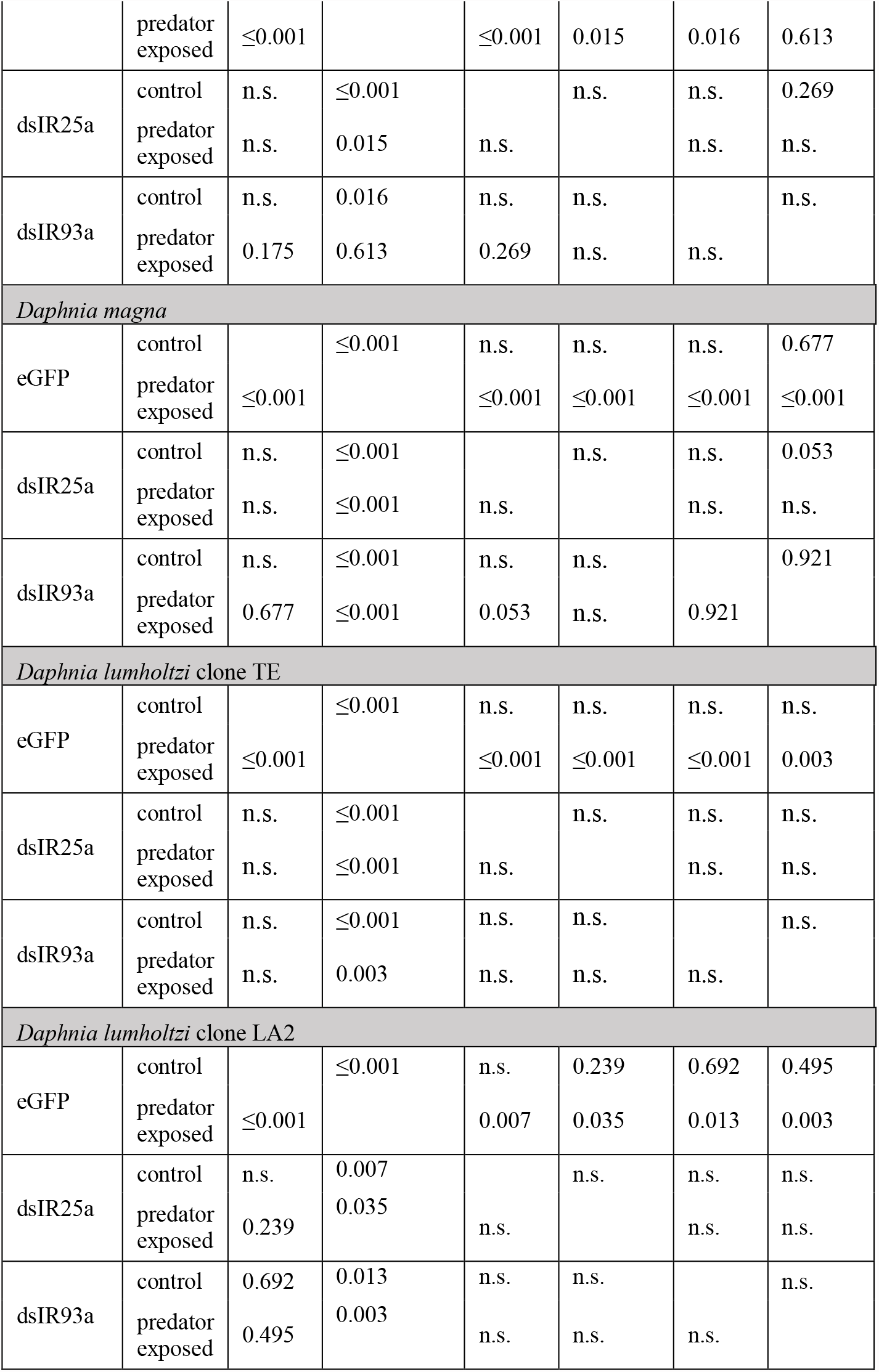
Post hoc Bonferroni analysis on log(normalized crest height) in *D. longicephala*, log(body width) in *D. magna* and log(normalized head spine length) in *D. lumholtzi* clones TE and LA2 after injection with eGFP-, dsIR25a-or dsIR93a-probe measured after 48 hours predator exposure or control treatment. Related to fig. 5.

**Table S3:**
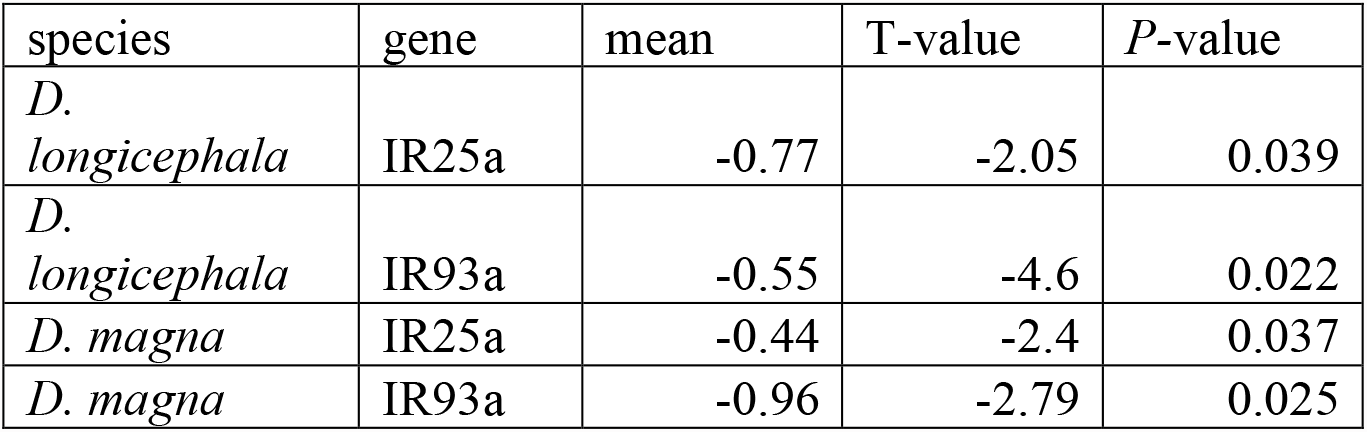
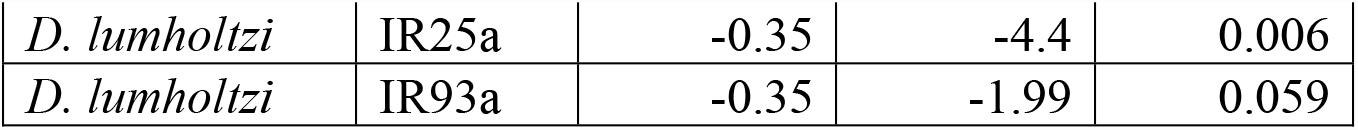
Differential gene expression statistical analysis results after probe injection as displayed in in fig. 5.

**Table S4:**
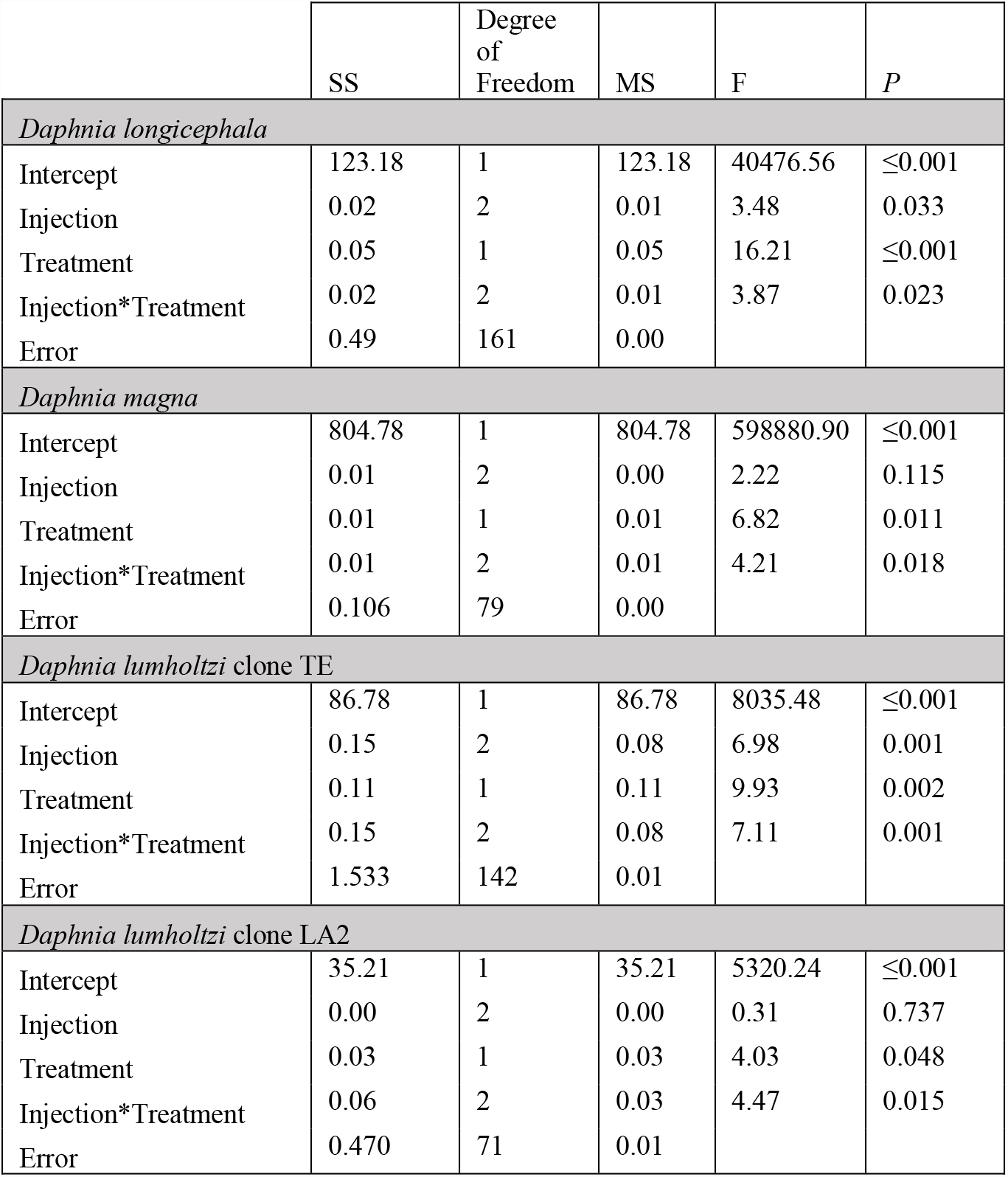
Factorial ANOVA on log(normalized crest height) in *D. longicephala*, log(body width) in *D. magna* and log(normalized head spine length) in *D. lumholtzi* clones TE and LA2 after injection with eGFP or the alternative IR25a or IR93a dsRNA probes measured after 48 hours predator exposure or control treatment. Related to fig. S5. Abbreviations: SS, square sum; MS, mean square

|  | SS | Degree of Freedom | MS | F | P |
| --- | --- | --- | --- | --- | --- |
| *Daphnia longicephala* |  |  |  |  |  |
| Intercept | 123.18 | 1 | 123.18 | 40476.56 | ≤0.001 |
| Injection | 0.02 | 2 | 0.01 | 3.48 | 0.033 |
| Treatment | 0.05 | 1 | 0.05 | 16.21 | ≤0.001 |
| Injection*Treatment | 0.02 | 2 | 0.01 | 3.87 | 0.023 |
| Error | 0.49 | 161 | 0.00 |  |  |
| *Daphnia magna* |  |  |  |  |  |
| Intercept | 804.78 | 1 | 804.78 | 598880.90 | ≤0.001 |
| Injection | 0.01 | 2 | 0.00 | 2.22 | 0.115 |
| Treatment | 0.01 | 1 | 0.01 | 6.82 | 0.011 |
| Injection*Treatment | 0.01 | 2 | 0.01 | 4.21 | 0.018 |
| Error | 0.106 | 79 | 0.00 |  |  |
| *Daphnia lumholtzi* clone TE |  |  |  |  |  |
| Intercept | 86.78 | 1 | 86.78 | 8035.48 | ≤0.001 |
| Injection | 0.15 | 2 | 0.08 | 6.98 | 0.001 |
| Treatment | 0.11 | 1 | 0.11 | 9.93 | 0.002 |
| Injection*Treatment | 0.15 | 2 | 0.08 | 7.11 | 0.001 |
| Error | 1.533 | 142 | 0.01 |  |  |
| *Daphnia lumholtzi* clone LA2 |  |  |  |  |  |
| Intercept | 35.21 | 1 | 35.21 | 5320.24 | ≤0.001 |
| Injection | 0.00 | 2 | 0.00 | 0.31 | 0.737 |
| Treatment | 0.03 | 1 | 0.03 | 4.03 | 0.048 |
| Injection*Treatment | 0.06 | 2 | 0.03 | 4.47 | 0.015 |
| Error | 0.470 | 71 | 0.01 |  |  |

**Table S5:**
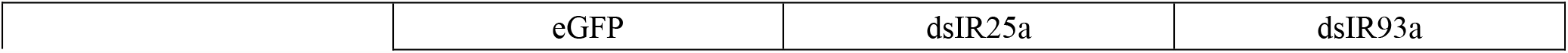
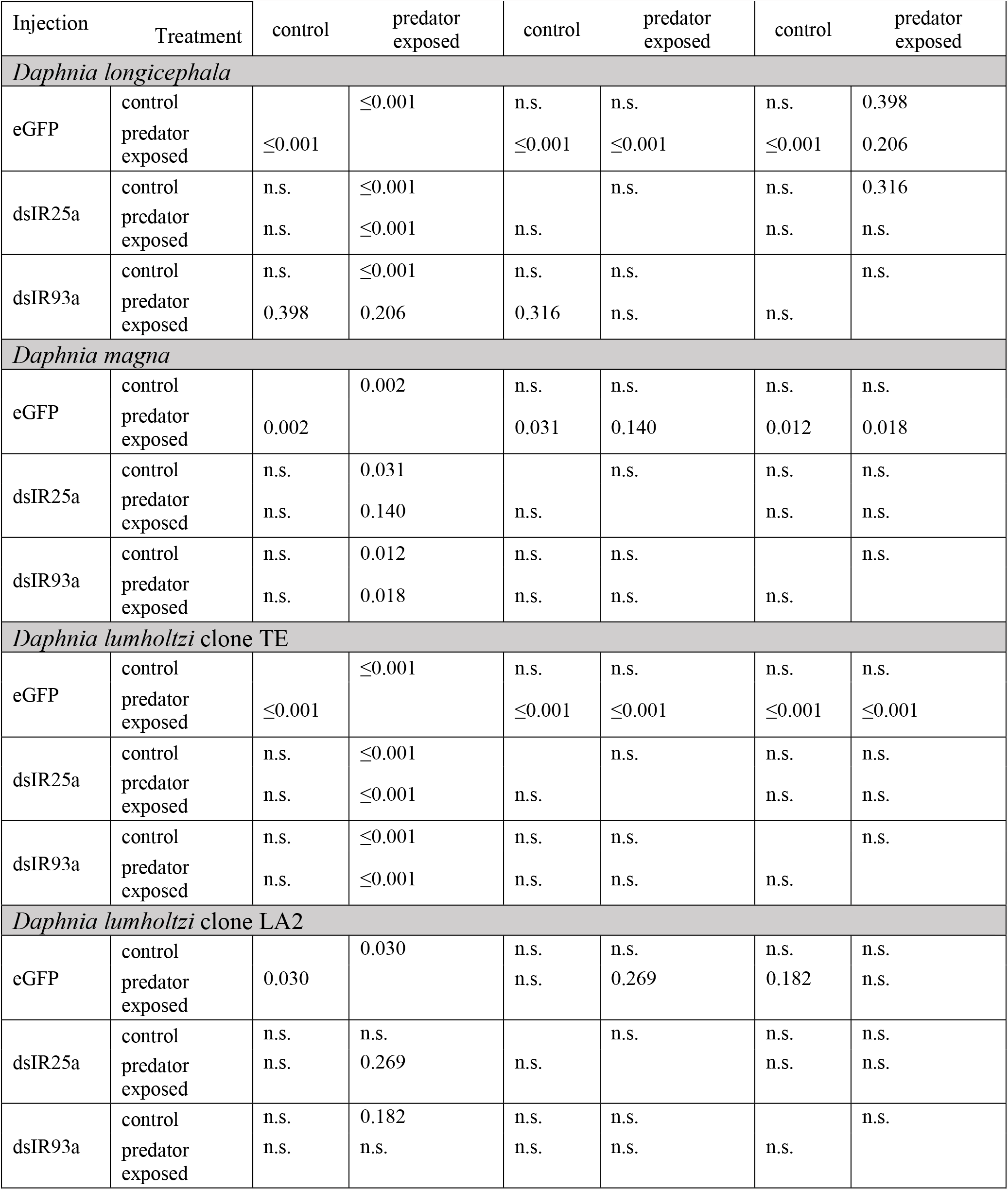
Post hoc Bonferroni analysis on log(normalized crest height) in *D. longicephala*, log(body width) in *D. magna* and log(normalized head spine length) in *D. lumholtzi* clones TE and LA2 after injection with eGFP-, dsIR25a-or dsIR93a-probe measured after 48 hours predator exposure or control treatment. Related to fig. S5.

